# Vimentin protects the structural integrity of the nucleus and suppresses nuclear damage caused by large deformations

**DOI:** 10.1101/566174

**Authors:** Alison E Patteson, Amir Vahabikashi, Katarzyna Pogoda, Stephen A Adam, Anne Goldman, Robert Goldman, Paul A Janmey

**Affiliations:** Institute for Medicine and Engineering, University of Pennsylvania, Philadelphia, PA 19104; Physics Department, Syracuse University, Syracuse, NY 13244; Department of Cell and Molecular Biology, Feinberg School of Medicine, Northwestern University, Chicago IL 60611; Institute of Nuclear Physics, Polish Academy of Sciences, PL-31342 Krakow, Poland; Department of Physiology, University of Pennsylvania, Philadelphia, PA 19104

## Abstract

Mammalian cells frequently migrate through tight spaces during normal embryogenesis, wound healing, diapedesis or in pathological situations such as metastasis. The nucleus has recently emerged as an important factor in regulating 3D cell migration. At the onset of migratory behavior, cells often initiate the expression of vimentin, an intermediate filament protein which forms networks extending from a juxtanuclear cage to the cell periphery. However, the role of vimentin intermediate filaments (VIFs) in regulating nuclear shape and mechanics remains unknown. Here, we used wild type and vimentin-null mouse embryonic fibroblasts to show that VIFs regulate nuclear shape, motility, and the ability of cells to resist large deformations. The results show that loss of VIFs alters nuclear shape, reduces perinuclear stiffness, and enhances motility in 3D. These changes increase nuclear rupture and activation of DNA damage repair mechanisms, which are rescued by exogenous re-expression of vimentin. Our findings show that VIFs provide mechanical support to protect the nucleus and genome during migration.

## Introduction

The proper function and homeostasis of tissues depends on the ability of individual cells to withstand demanding physical stresses. For example, during migration, a cell must squeeze through small interstitial spaces in tissues imposing large strains on entire cell bodies and their largest organelle, the nucleus. Determining how cells maintain their structural integrity under these large strains is an important pre-requisite for understanding a wide range of normal physiological activities including tissue morphogenesis during development, wound healing, diapedesis, and pathological conditions such as cancer cell metastasis and chronic inflammatory diseases such as arthritis.

The deformability of cells depends largely on the cytoskeleton, which is comprised of three main polymers, F-actin, microtubules, and intermediate filaments (IFs). F-actin and microtubules are highly conserved in eukaryotic cells and single-celled organisms, but IFs are diverse and evolved later as multicellular organisms appeared. Vimentin is a Type III IF protein, and has been used extensively as a reliable marker of the epithelial to mesenchymal transition (EMT), in which non-migratory epithelial cells lose cell-cell adhesions, dramatically alter their shape, and transition to a highly migratory mesenchymal phenotype ^1–4^. Furthermore, vimentin intermediate filaments (VIFs) are implicated in the development of multiple cancers, and the expression of vimentin is a clinical marker of poor prognosis and increased metastasis ^5^. Yet, little is known regarding the role of VIF in 3D cell motility.

One robust feature of the organization of VIFs is their assembly into an intricate cage-like network that surrounds the nucleus ^6^. The juxtanuclear VIF network extends throughout the cytoplasm to the cell surface helping to position the nucleus ^7, 8^ and other organelles ^9, 10^. Even under conditions in which the peripheral VIF network is dynamic, such as during growth on soft substrates, the perinuclear VIF cage remains intact ^11^. There is evidence that VIFs establish indirect physical connections to the outer nuclear membrane through interactions with the linker of the nucleoskeleton and cytoskeleton (LINC) complex ^12^. The LINC complex has also been shown to connect to the nuclear lamina, a thin filamentous layer surrounding the nuclear periphery that is mainly composed of the type V IF proteins, the nuclear lamins ^13, 14^. There is considerable evidence that the nuclear lamina plays an important role in determining nuclear shape and rigidity ^15–19^. Changing the expression patterns of specific lamin isoforms alters nuclear shape and the lamin meshwork structure ^15, 20, 21^ and can lead to nuclear abnormalities, such as blebs, in which the lamin B isoform is depleted ^21^. Nuclear blebs can also occur spontaneously during cell migration through confined spaces ^22, 23^. Blebbing can lead to rupture of the nuclear envelope, unregulated mixing of the nuclear and cytosolic materials, accumulated DNA damage, and genomic instability ^24^.

Given the proximity of VIFs to the nucleus and their connections to the cytoplasmic face of the nuclear envelope, it is likely that they also play a role in regulating the mechanical properties of the nucleus. This idea is supported by the findings that VIF are viscoelastic biopolymers that stiffen at large strains and unlike crosslinked actin or microtubules are capable of withstanding extreme deformations without breakage ^25, 26^. Based upon these properties, our central hypothesis is that the perinuclear cage of VIF provides a protective structure that resists extreme deformations of the nucleus and thereby protects the structural integrity of the cell. Here, we investigate the effects of VIF on nuclear morphology and stability in cells grown on conventional 2D rigid substrates and in 3D confining environments, where squeezing of the cells through small pores can lead to large nuclear deformations and damage ^22, 23, 24^.

## Results

### The perinuclear VIF cage regulates nuclear shape and volume

We used confocal microscopy to examine the structure of the VIF network around the nucleus in vim +/+ mouse embryonic fibroblasts (mEFs). The confocal stacks captured at the basal (Fig. 1a) and apical (Fig. 1b) nuclear surfaces show the proximity between the VIFs and the nuclear surface. Furthermore, the 3D renderings of the confocal stacks also show that VIFs are closely juxtaposed to the nuclear surface, frequently appearing as a cage surrounding it (Fig. 1 c). Details of the vimentin cage structure at the nuclear envelope are shown in Structured Illumination Microscope (SIM) images of a WT mEF at the basal (Fig. 1d) and apical surface (Fig. 1e), and also maximum projection of the z-stacks (Fig 1f) of the vimentin network.

**Fig. 1.**
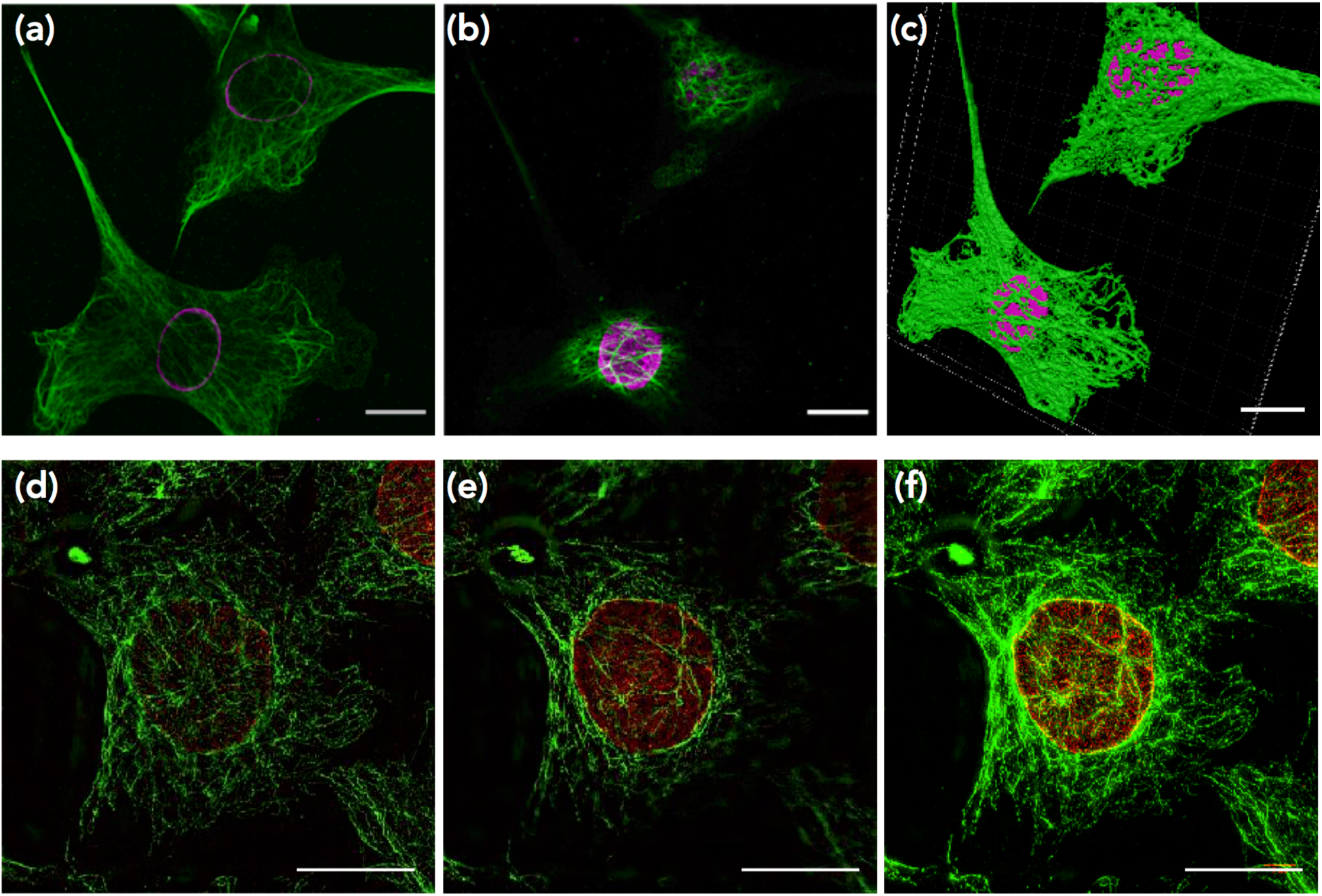
Perinuclear cage of VIF. Confocal images of the VIF (green) cage at the lower (a) and upper (b) surfaces of the nuclei (magenta) and their 3D rendering (c) in two WT mEFs. SIM images of the vimentin cage around the nucleus (red) in a vim +/+ mEF at the basal (d) and apical plane (e). (f) is the maximum projection of SIM images for the cell in (d) and (e). Scale bar is 20 μm.

Figure 2 shows that the robust vimentin network in vim +/+ mEFs (Fig. 2a) is missing in vim −/− mEFs (Fig. 2b). Expression of exogenous wild-type vimentin in vim −/− mEFs (which we refer to as “rescued mEFs”) results in variable expression levels of vimentin in these cells, averaging ∼20% of that seen in vim +/+ cells as determined by immunoblotting. However, this level of vimentin expression in rescued mEFs is sufficient to form a partial nuclear cage with some associated filamentous structures extending towards the cell surface (Figs. 2c-d).

**Fig. 2.**
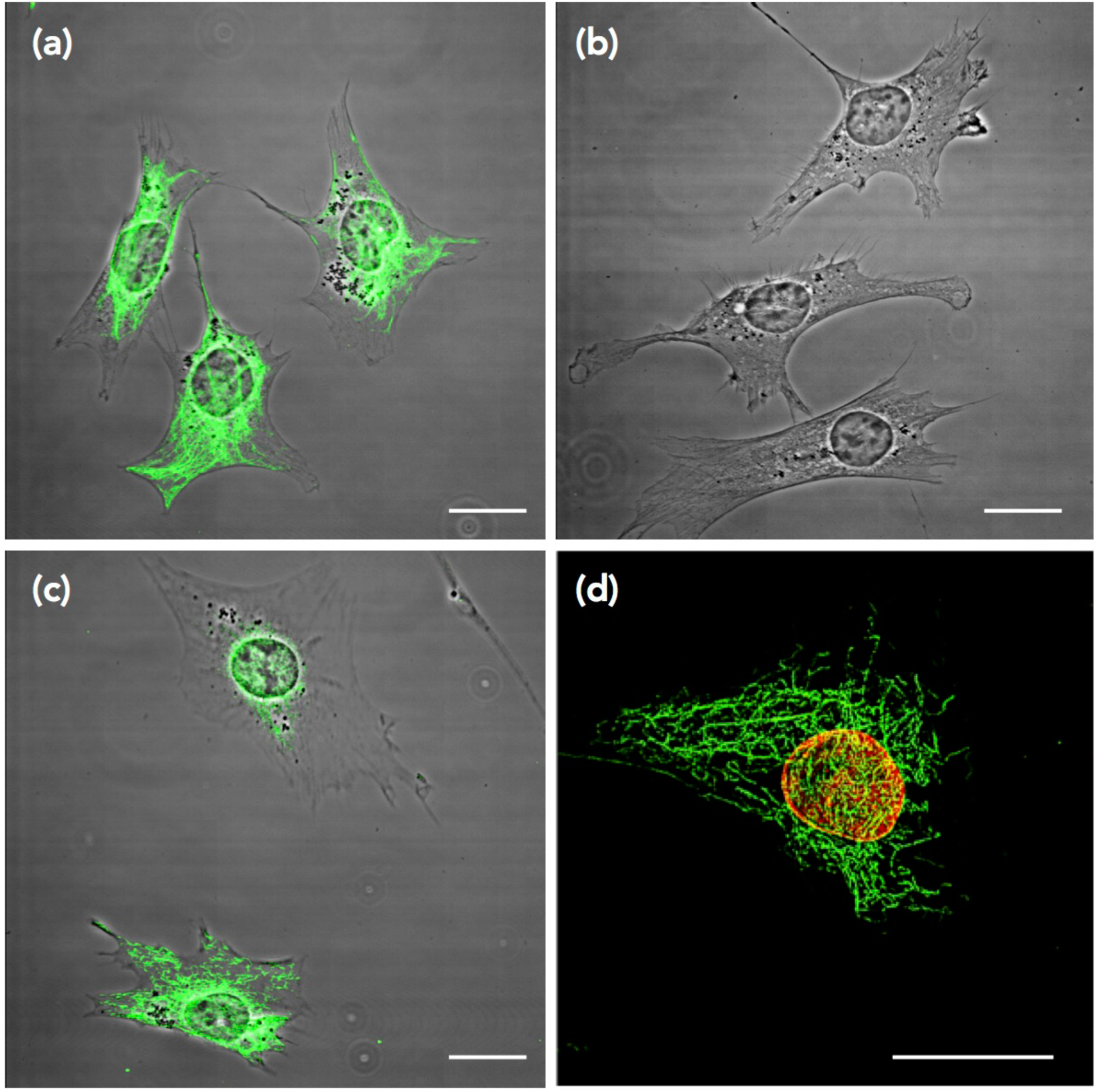
Overall Distribution of VIF in vim +/+, vim−/−, and rescued mEFs. Immunofluorescence (maximum intensity projection of confocal stacks) / phase contrast images of vimentin (green) in vim +/+ (a), vim −/− (b), and rescued mEFs expressing exogenous wild-type vimentin (c). SIM image of the vimentin cage around the nucleus (red) in a rescued mEF (d). Scale bar is 20 μm.

Given the proximity of the VIF cage to the nuclear surface and the known strain stiffening properties of VIF, we determined whether it is involved in regulating nuclear shape. To this end, we compared nuclear profiles in vim +/+ and vim −/− mEFs. The nuclei in vim +/+ mEFs adhering to a flat rigid surface typically possess an oblate spheroidal shape. In contrast, the nuclei in vim −/− mEFs show an altered shape, looking significantly rounder compared to the vim +/+ mEFs (Fig. 3a). Expressing exogenous wild-type vimentin in the vim −/− mEFs did not rescue the average nuclear shape phenotype. This is likely attributable to the low vimentin expression levels as reflected by the incomplete restoration of the VIF cage in the rescued cells. To quantify the changes in nuclear shape, we calculated the sphericity of nuclei in vim +/+, vim −/−, and rescued mEFs. The sphericity analysis showed that nuclei in vim +/+ mEFs are significantly flatter and more elliptical in shape when compared to the vim −/− and rescued mEFs that possess rounder nuclei (Fig. 3b, p<0.0001). We next used confocal stacks to generate 3D renderings of the nuclei to calculate their corresponding volume in vim +/+, vim −/−, and rescued mEFs. The average nuclear volumes in the vim −/− and rescued mEFs were smaller by over 25% when compared to the vim +/+ mEFs (p<0.0001, Fig. 3c). There was no difference between the vim −/− and rescued cells (p=0.20). Similar results were obtained for cell volume where the average volume of vim +/+ mEFs was greater than that of the vim −/− and rescued mEFs (p<0.025, Fig. 3d). Vim −/− and rescued mEFs showed comparable average cell volumes (p=0.9). We also found a significant correlation between the nucleus and cell volume in all cell types examined (p<0.002, Fig. 3e). However, the regression slope for the vim −/− mEFs (6.9) was significantly different from those for vim +/+ (9.5) and rescued mEFs (9.0) (p<0.04, SI Fig. 1), suggesting that for the same total cell volume, the nuclei of vim −/− cells tend to be smaller.

**Fig. 3.**
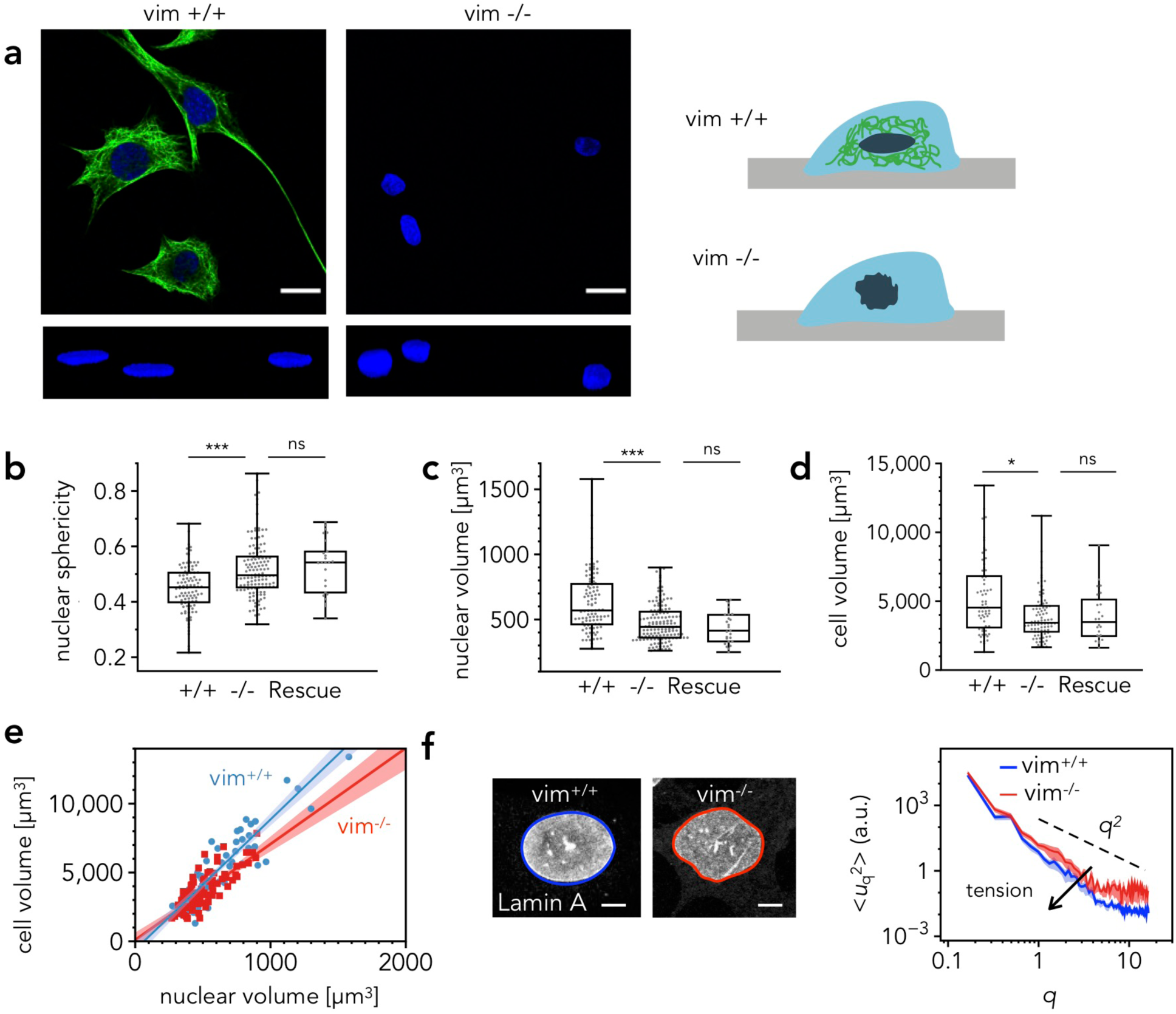
VIF cage regulates nuclear shape and volume. Confocal image of the vimentin network (top, green) and nucleus (blue), and 3D renderings of the nuclei (bottom) in vim +/+ and vim −/− mEFs (a); Images show loss of vimentin changes nuclear morphology. Scale bar is 20 μm. The nuclei of cells lacking vimentin are rounder and less smooth compared to those in vim +/+ mEFs. Vim +/+ mEFs have more compressed, elongated nuclear shapes compared to the vim −/− and rescued mEFs expressing relatively low levels of vimentin (b). Lack of vimentin decreases nuclear (c) and cell volume (d). Nuclear volume correlates with cell volume in vim +/+, vim −/−, and rescued mEFs (e). Quantification of the nuclear structure by the Fourier amplitudes of the nuclear shape, u(q) (n = 40-43) in fluorescent images of lamin A in vim +/+ and vim −/− mEFs (f). Lamin A immunofluorescence was used to trace nuclear contour. The normalized amplitude decreases as q^2^ with a prefactor based on undulated membrane theory that is inversely proportional to the nuclear stress. Scale bar is 5 μm.

Lastly, we found that the surface of the nucleus was smoother in vim +/+ mEFs compared to the vim −/− ones, as shown by lamin A staining in Fig. 3f. These results are consistent with prior studies using SW-13 cells, which showed that in cells without vimentin the nuclear morphology was more folded and less smooth compared to cells with vimentin ^27^. To quantify the structure of the nucleus, we traced the nuclear contour r(θ) and calculated the mean-square amplitude of the Fourier modes of nuclear shape 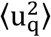 as described in Methods. The mean squared amplitude decays with wave number q, scaling as 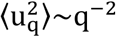, which is similar to tension-suppressed fluctuation in lipid membranes ^28^, a model that has previously been applied to cell nuclei ^29^. According to such a model, the suppression of shape fluctuations in the presence of VIF corresponds to an increase in nuclear tension or stress.

### VIFs confer mechanical stability to the nucleus

To assay the mechanical stability of the nuclear envelope (NE), we imaged the incidence of nuclear rupture in mEFs transfected with GFP coupled to a nuclear localization signal (NLS-GFP). Live cell imaging confirmed that the NLS-GFP accumulated in the nucleus (Fig. 4a,c). Therefore the loss of NLS-GFP fluorescence intensity from the nucleus and into the cytoplasm was indicative of NE rupture events ^20, 22, 30^ (Fig. 4a,c). In a 17 hr observation window, ruptures occurred in 8% of vim+/+ and 18% of vim −/− mEFs cultured on 2D substrates, respectively (Fig. 4b). The NE ruptures seemed to correlate with cells containing blebbed nuclei, as reflected by increase in cytoplasmic fluorescence intensity. Nuclear envelope rupture was more common in 3D collagen gels when compared to the 2D substrates (Fig. 4 b,d, *p*<0.01 for vim +/+ and vim −/− mEFs). Cells cultured in collagen gels exhibited more nuclear blebs and ruptures. These were most frequently seen in cells that migrated through a small pore that constricted the nucleus to a diameter of 2 – 3 µm. After rupture many nuclei were repaired, presumably by the endosomal sorting complex, and the NLS-GFP was transported back into the nucleus ^22, 23^. The incidence of NE rupture and repair was more frequent in vim −/− mEFs (Fig. 4b, d; SI Movies 1&2); specifically, in the 3D collagen gel assays, the percentage of ruptured nuclei in vim −/− mEFs was more than 70% higher compared to the vim +/+ mEFs (Fig. 4d; p = 0.03).

**Fig. 4.**
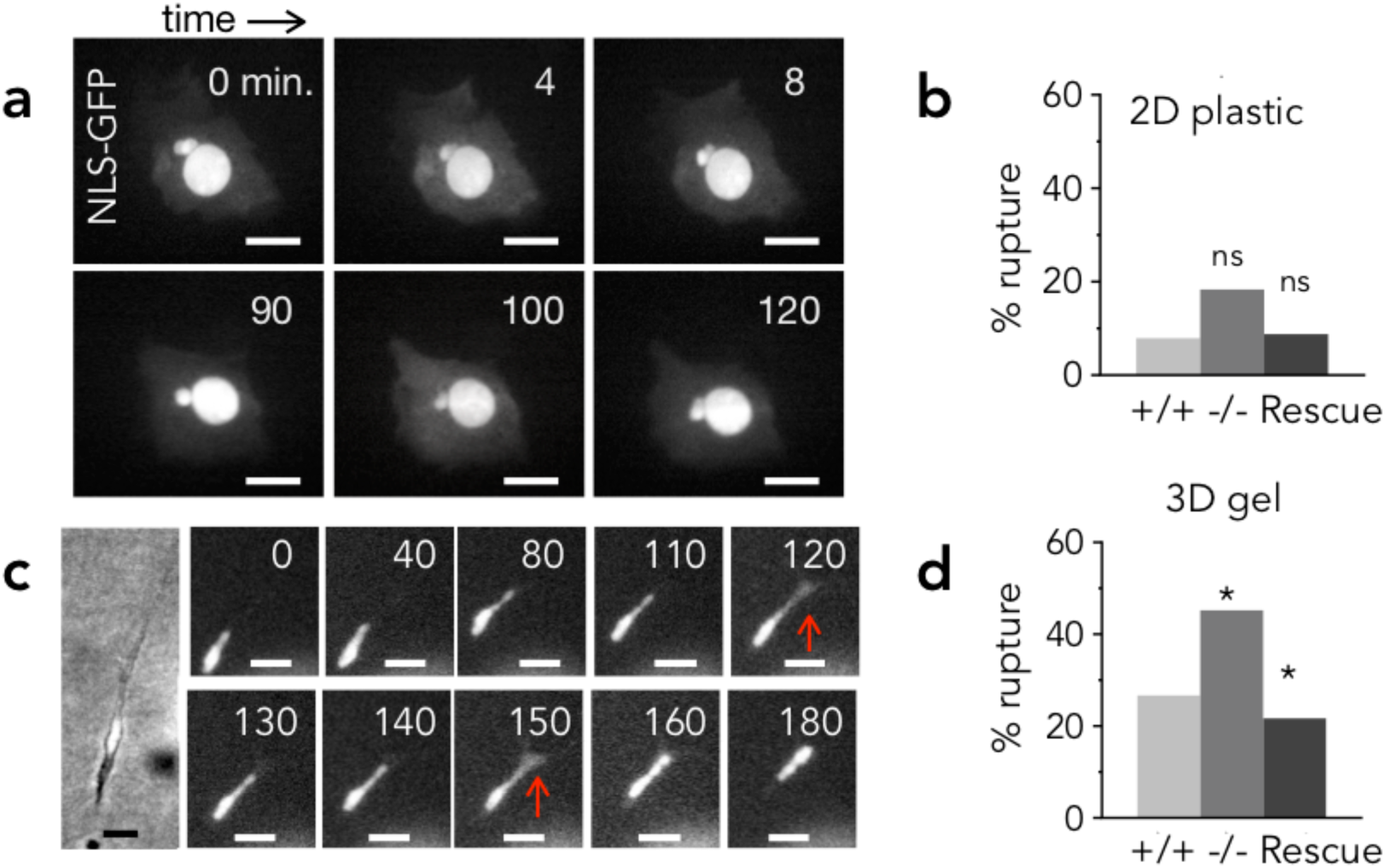
Loss of VIF increases nuclear envelope rupture. Sequential images of vim −/− mEF expressing NLS-GFP on 2D tissue culture plastic (a) and in 3D collagen gels (c). Leakage of NLS-GFP signal from the nucleus to the cytoplasm indicates nuclear envelope ruptures. Scale bar is 20 μm. Fraction of cells with nuclear membrane rupture in 2D (b) and 3D (d) environments (as observed over a 17 hr interval). Vimentin reduces nuclear membrane ruptures and thereby promotes the mechanical stability of the nucleus, particularly in confining 3D gels. The vim −/− phenotype is rescued by the expression of vimentin. (In (a), n = 57-63 cells per condition, and in (c), n = 37-53 cells; data collected over a minimum of two experiments). Denotation *, p < 0.05.

Exogenous expression of wild-type vimentin in vim −/− mEFs restored the nuclear rupture rate to that of vim +/+ cells in both 2D and 3D assays, despite the fact that the average level of vimentin expression in rescued cells was ∼ 20% of the vim +/+ mEFs (Fig. 4b, d). This indicates that a relatively small number of perinuclear VIF can rescue the nuclear envelope rupture phenotype in vimentin −/− mEFs (see Fig 1).

### VIFs protect against extreme nuclear deformation and damage during migration through small pores

To examine the role of VIF in nuclear deformations induced by 3D cell migration, we seeded cells on rigid filter membranes and examined their motility (Fig. 5a). Results from the migration assays show that vim −/− mEFs transit more readily through the pores of all three diameters (3,5, and 8 µm) compared to the vim +/+ cells (p<0.01), and that the expression of vimentin in the vim −/− cells rescues the wild-type migration rates (Fig. 5a). The results differ from those on 2D surfaces where removal or disruption of the VIF network in fibroblasts reduces motility ^31, 32^. Here, the faster movement of vimentin-null cells through the pores implies that additional work is needed to deform the vimentin network in normal cells as they migrate through the 3D constriction. Results from immunofluorescence studies show that VIF remain associated with the nucleus after cells translocate through the pores (Fig. 5b, image shown for a 5 µm pore), suggesting that the VIF network can withstand large strains to protect the nucleus during translocation. In addition, the nuclei in cells that migrated through the membrane had a significantly smaller area (as determined by manually tracing 2D confocal images of Hoechst-stained nuclei on the membrane) compared to those on the top, for both vim +/+ and vim −/− mEFs (p<0.01, Fig. 5c). This finding indicates that the small pore size of the membrane selects for the passage of cells with smaller nuclei, consistent with the nucleus acting as a rate-limiting step through constricted migration ^33–35^.

**Fig. 5.**
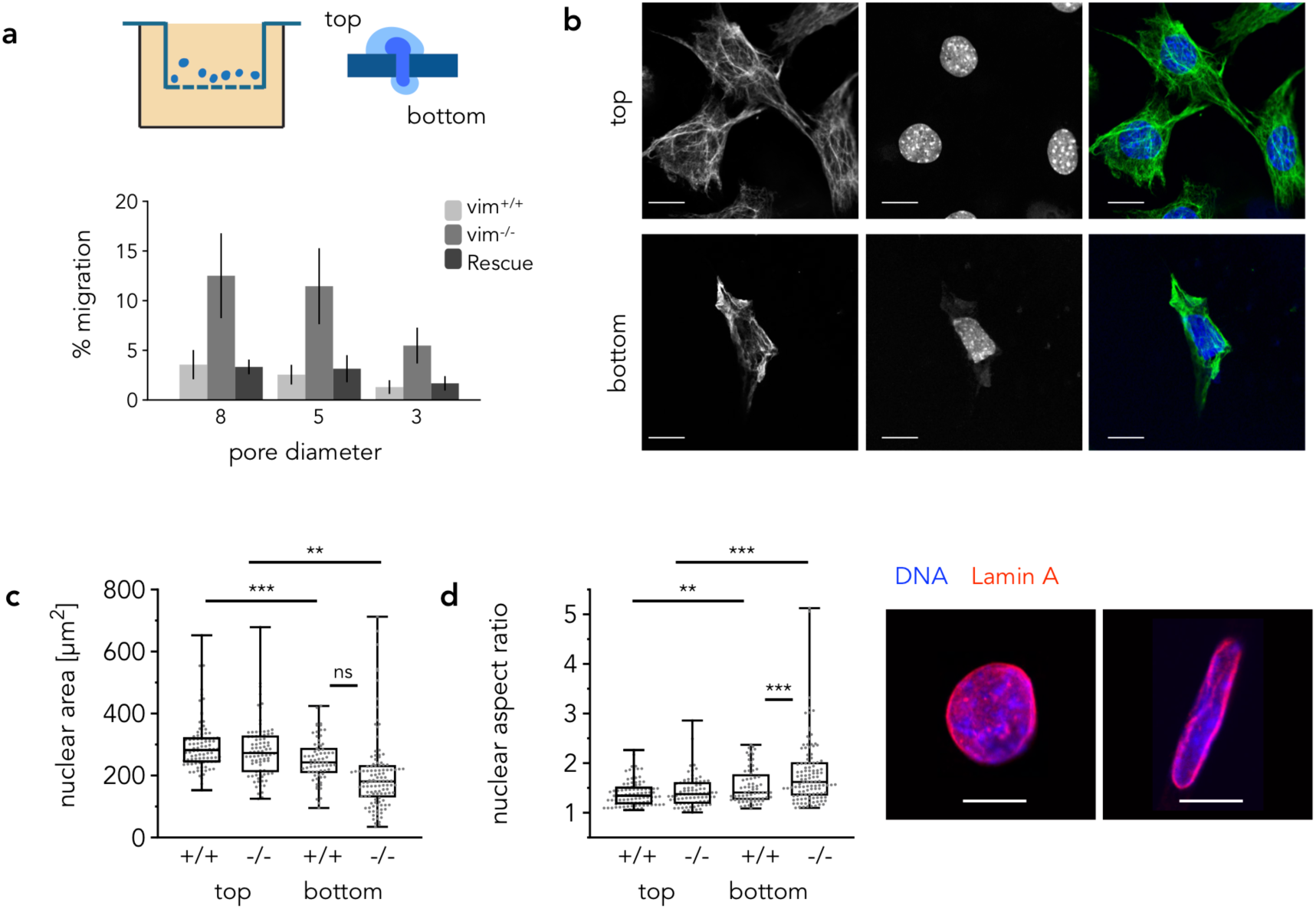
VIF hinders migration through small pores but limits nuclear deformation. Cells are seeded on Transwell membranes and allowed to migrate through pores in the membrane (see schematic). Loss of vimentin increases migration through membranes of varying pore sizes and normal cell motility is restored upon vimentin re-expression (a). Images of vim +/+ cells fixed and stained for vimentin and the nucleus (Hoechst) on the top and bottom of the membrane. Scale bar is 20 μm. The membranes select for smaller nuclei, as shown by the decrease in nuclear area for cells on top and bottom of the membrane (c). Pore migration increases nuclear aspect ratio (d); In cells lacking vimentin, nuclei are stretched to a greater extent. Immunostaining shows nuclear morphologies on the bottom of the membrane in vim +/+ and vim −/− mEFs. Scale bar is 10 μm.

Migration through small pores can cause nuclear shape changes that depend on the mechanical properties of both the nucleus and whole cell ^35^. Based on this finding, we determined whether nuclear shape in the vim +/+ and vim −/− mEFs was altered following their migration through 3 µm pores (Fig. 5d). We found that nuclear shape was altered in both vim +/+ and vim −/− mEFs after migration (p<0.01), and this effect was greater for cells lacking vimentin (p<0.001).

The mEFs, particularly vim −/− ones, exhibit nuclear blebbing after migrating through the 3 µm pores. Immunofluorescence revealed that post-migration nuclear blebs contained mainly dilated lamin A meshworks and little if any lamin B (Fig. 6a,b,c). These lamin A rich blebs devoid of lamin B have been reported in a variety of cell types and in different experimental conditions ^21, 22^. Translocating through the pores increased the fraction of nuclei with blebs in both cell types (Fig. 6d, p<0.001) with vim −/− mEFs containing approximately 35% more nuclei with blebs compared to the vim +/+ mEFs (p=0.03).

**Fig. 6.**
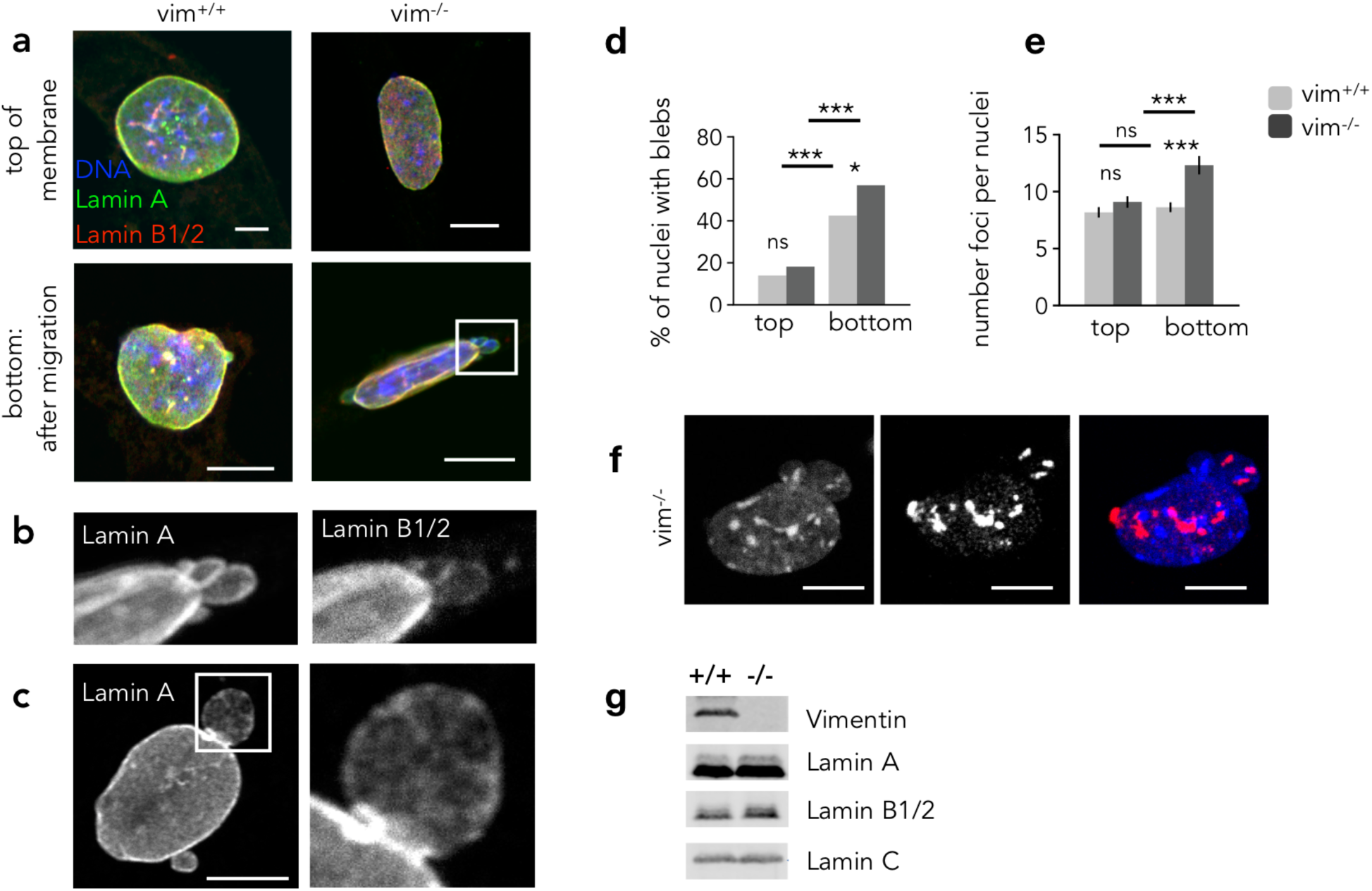
Loss of vimentin increases nuclear damage but does not alter lamin expression levels. Immunostaining images showing lamin A and lamin B1/2 on the top and bottom of 3 μm membranes with 3 μm pores (a); On the membrane bottom, vimentin-null mEF exhibit nuclear abnormalities such as large nuclear blebs (b) and some blebs exhibit dilated lamin A networks (c). Loss of vimentin increases the percentage of nuclei with migration-induced blebs (d). Loss of vimentin increases the number of post-migration DNA breaks, quantified by the number of ƔH2AX foci per nucleus (e); Immunostaining images show ƔH2AX foci, a marker of double-strand DNA breaks, in vim −/− mEFs nuclei on the membrane bottom (f): foci appear throughout the nucleus and in protruding blebs. vim +/+ and vim −/− mEFs express equivalent levels of lamin A, B1/2, and C as shown by quantitative immunoblotting (g). Scale bar is 10 μm.

It has also been shown that extreme mechanical deformation of the nucleus during cell migration results in increased DNA damage ^24, 36^, and that nuclear ruptures are related to the sustained loss of DNA damage repair factors ^22, 23^. Therefore, we determined whether loss of VIF is also associated with increased DNA damage during constricted migration (Fig. 6e and f). DNA damage repair was assayed by immuno-staining for Ɣ-H2AX, a marker of double-strand DNA breaks ^37^. The results show a 50% increase in migration-induced DNA damage repair foci in vim −/− mEFs (p<0.001) as compared to the vim +/+ mEFs. This is consistent with the increased NE rupture data shown in Figure 4.

There is abundant evidence that the mechanical integrity of the nucleus, or even the whole cell, depends on the expression levels of nuclear lamin A which along with the B Type lamins form the main structural component of the nuclear lamina underlying the nuclear envelope ^15, 16, 18^. We therefore determined the lamin expression levels in vim +/+ and vim −/− mEFs by quantitative immunoblotting. Figure 6g shows that vim +/+ and vim −/− mEFs have indistinguishable amounts of lamins A, B1/2, and C, verifying that lamin isoform expression levels are not altered in vim −/− mEFs. Thus, the changes detected in our experiments reflect the role of VIFs in nuclear mechanics, independent of effects from nuclear lamins.

### Vimentin deficient cells have compromised perinuclear mechanics

Previous studies have shown that vim −/− mEFs have compromised cortical and cytoplasmic stiffness in peripheral regions away from the cell nucleus ^38, 39^. In this study we measured the cytoplasmic stiffness of cells immediately above the nucleus by atomic force microscopy (AFM) using large AFM round tips on the AFM cantilever (Fig. 7a, b). The AFM measurements show that the apparent Young’s modulus over the nucleus is more than 30% lower in vim −/− mEFs. This reduction in Young’s modulus is partially reversed in the vimentin rescued cells (Fig. 7b). These results show that under compressive loads the lack of VIF leads to greater cell deformation and, potentially, greater damage to the cell and nuclear envelope.

**Fig. 7.**
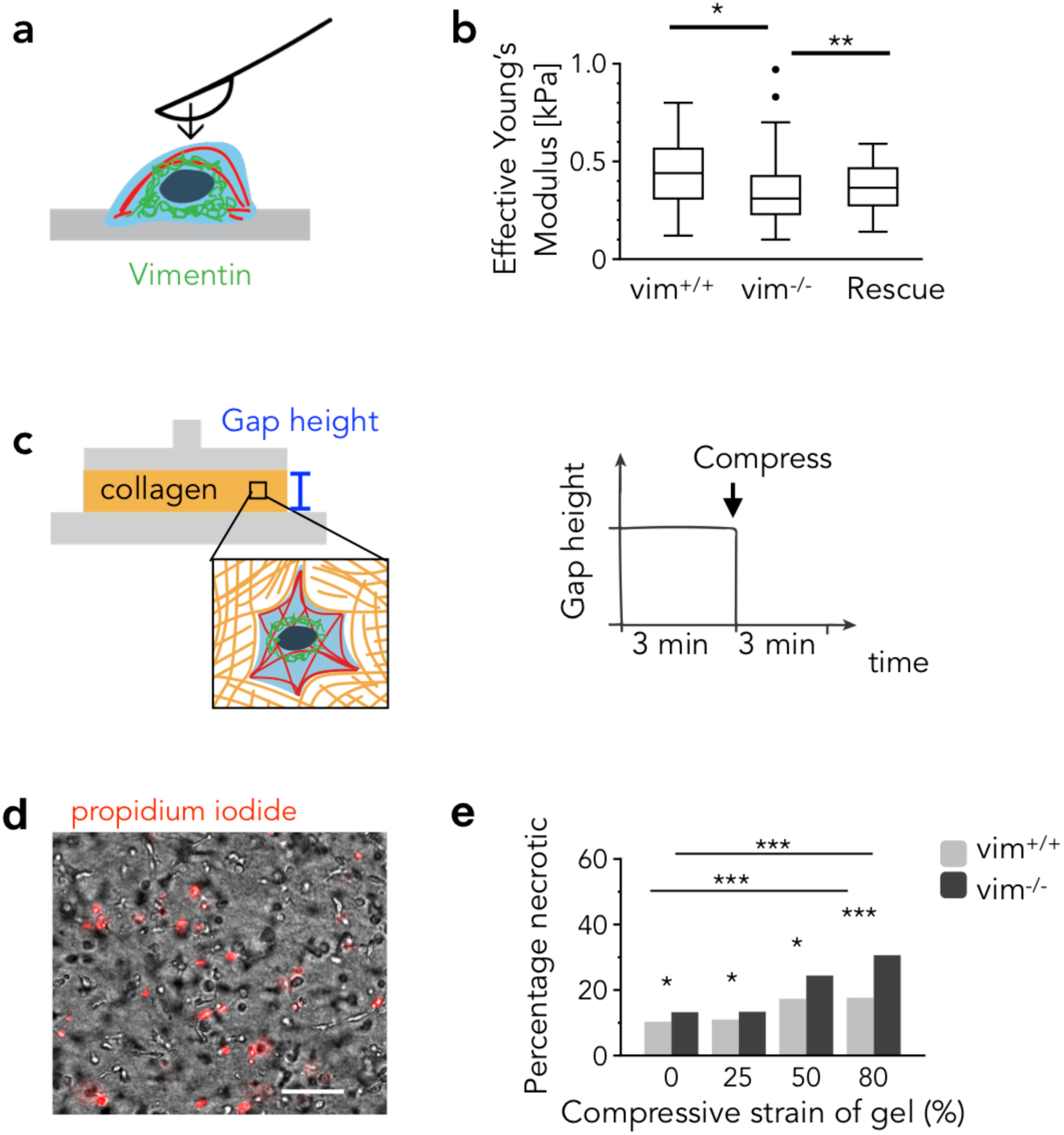
VIF provides mechanical resistance and protection against compressive forces. Schematic for the AFM measurements on vim +/+, vim −/−, and rescued mEFs (a). The average Young’s modulus of the vim −/− mEFs is lower than vim +/+ mEFs (*p*<0.04) and rescued mEFs (p<0.01) (b). Schematic image for external compression of a model system of mEFs cultured in 3D collagen gels (c); The gels are compressed with a parallel plate rheometer at varying degrees of axial strain. Cells are stained with propidium iodide to determine the amount of necrotic cell death, as shown for 80% compression (d). Scale bar is 100 μm. Vim −/− mEFs exhibit more necrosis at large strains compared to the vim +/+ mEFs, indicating that VIF protect the structural integrity of the cell by resisting large compressive strains (e).

### VIF reduce necrosis resulting from large external compressions

VIFs are expressed in cells of tissues that are routinely subjected to mechanical stresses ^40–42^. Examples include the retina and fat tissues, which regularly undergo compression and extension, and tumors, which generate solid stress ^43^. Whether or not vimentin protects individual cells from damage due to large strains has yet to be determined. We tested the relation between vimentin expression, externally applied strains, and cell fate *in vitro*. We cultured vim +/+ and vim −/− mEFs in 3D collagen gels and subjected the gels to axial compression with a parallel-plate rheometer (Fig. 7c). Cell death was detected by positive staining with propidium iodide, a marker of ruptured cell membranes and necrosis (Fig. 7d).

In the absence of external compression (0% strain), both vim +/+ and vim−/− mEFs had basal levels of necrotic cell death, presumably resulting from the stresses that the contractile cells themselves apply to the matrix and to restrictions to solute diffusion at the gel boundaries. Necrosis in vim −/− mEFs was 30% higher than wild-type’s in uncompressed gels (p=0.01, Fig. 7e), which could stem from the motility-induced damage and nuclear envelope rupture as shown in Figure 4. This basal level was not due to confinement of the cell-containing gel between the metal plates of the rheometer because the levels were similar for uncompressed gels in a culture dish and for minimally compressed cells in the rheometer. Large axial compressions of the cell-seeded collagen matrices led to necrosis in both cell types; however, for the large compressive strains (80%), vim −/− cells had a 170% increase in necrosis compared to their wild-type counterparts (p<0.001, Fig. 7e). The compressive stresses to produce these levels of strain ranged from 0-100 Pa. Since the Young’s moduli of the cell-containing gels at small strains (3x shear modulus ∼ 90-210 Pa from SI Fig. 2 by parallel plate rheology) were smaller than that of the cells (approximately 300-400 Pa from Fig 6c by AFM) the individual cell strain was correspondingly lower than that of the whole gel, suggesting that significant necrosis occurs in the vim −/− mEFs under physiologically reasonable levels of compressive stress and strain. Overall, these results highlight that vimentin protects the cell from both passive and active deformations that can cause damage and control cell fate.

## Discussion

Mechanically induced deformations of the nucleus in cells migrating through small channels can cause nuclear envelope blebbing, rupture, and DNA damage. To date these changes have mainly been attributed to regulation by the nuclear lamin A, one of 4 lamin isoforms belonging to the Type V intermediate filament protein family ^22, 24, 44–46^. The lamins are major filamentous components of the nuclear lamina, which is closely associated with the inner membrane of the nuclear envelope ^46, 47^. The role of lamins in nuclear mechanics, especially under conditions of high deformation, is supported to some extent by the strain stiffening properties of the lamina ^16, 35, 48^. Recent studies utilizing cryo-EM tomography have revealed that the lamina is only ∼14 nm thick and that lamin filaments cover ∼50% of the nuclear surface ^49^. It is unlikely that this very thin layer of lamins at the nuclear surface is sufficient to protect the nuclear contents from the large mechanical stresses that occur as cells move through tight spaces or are compressed by adjacent tissues. It is for this reason that we have undertaken a study of the role of the Type III vimentin intermediate filaments forming the juxtanuclear cage. This is important as the strain stiffening properties of VIFs and their resistance to large deformations have been well documented ^26, 50^.

Our results show that in the absence of vimentin, cells migrate more frequently through small spaces, suggesting that perinuclear vimentin cushions the nucleus during the extreme strains associated with movement through constrictions. These results also suggest that VIFs function to maintain nuclear shape against the imposed deformations, acting like an elastic spring, which is also consistent with the viscoelastic and strain stiffening properties of VIFs ^25^. Furthermore, low levels of vimentin expression in vim −/− mEFs sufficient to partially restore the VIF cage effectively improved the mechanical stability of the nucleus and reduced nuclear envelope rupture, even though such low levels of expression failed to restore normal nuclear shape and the perinuclear stiffness.

Previous studies have demonstrated a role for the actin cytoskeleton in nuclear shape where the flat oblate nuclear profile has been attributed to the compression forces from the acto-myosin machinery ^51, 52^. However, in this study we find that vim −/− mEFs have altered nuclear shapes and are significantly rounder compared to the vim +/+ mEFs even though we detect no significant differences in F-actin distribution between these two cell types (SI Fig. 3). Furthermore, previous immunoblotting studies have shown that actin levels are the same in vim +/+ and vim −/− mEFs ^9^. The shape fluctuation analysis described in our studies suggests a decrease in nuclear tension in vim −/− mEFs that could also contribute to the nuclear rounding observed in these cells.

A potential explanation for the role of juxtanuclear VIF in regulating nuclear shape is its known interactions with F-actin via cytolinkers such as plectin, which also bind VIFs ^43, 44^. Alternatively VIFs may regulate nuclear shape by exerting tension on the nucleus through their connections to Nesprin-3, a member of the LINC complex, which interacts with both plectin ^53^ and actin ^54^. Recent studies indicate that during cell migration through extracellular matrices, the nucleus compartmentalizes the cell and is pulled towards the leading edge by actomyosin filaments. These latter filaments act through VIFs and nesprin-3, to create the high pressure required for lobopodia formation at the leading edge of locomoting cells ^55^.

Based upon our results we expect that cells subjected to high levels of mechanical stress may enhance vimentin expression in order to protect the nucleus against large strains but still allow for migration through tissues. In support of this, it has recently been shown that VIFs regulate both the mechanical properties of the cell cortex, the subcortical cytoplasm and the generation of traction forces required for cell motility ^39^.

The lower apparent tension of the nucleus of vim −/− cells at rest, inferred from their rounder shape (Fig. 3a) and higher degree of fluctuation (Fig. 3f) might appear at odds with the finding that the VIF perinuclear cage protects the nucleus from rupture. Physically, the explanation lies in the highly non-linear response of VIF networks to applied stresses. At low stress and strain, VIF networks are soft, and therefore even if they transmit forces exerted by the cortical cytoskeleton when cells adhere to a rigid surface, they effectively average out these force, preventing the large curvature deformations that can lead to nuclear envelope rupture ^56^. However, at large deformations, the VIF networks become much stiffer, and therefore can take up the stress generated by the cytoskeletal contraction that drives the cell through small spaces, sparing the nucleus from these large, transient stresses.

Taken together our results demonstrate a new role for VIFs in maintaining nuclear shape and mechanical stability. We establish that VIFs regulate the shape of the nucleus and that during migration through small channels their absence causes nuclei to undergo higher degrees of deformation, blebbing, and DNA damage in response to compressive forces. Our results provide new insights into the protective role of VIF in cells undergoing large strains with implications for cell fate, genome expression, healthy tissue maintenance, development, and disease progression.

## Methods

### 1. Cell culture

Vim +/+ and vim −/− mEFs were kindly provided by J. Eriksson (Abo Akademi University, Turku, Finland) and maintained in DMEM with 25 mM HEPES and sodium pyruvate (Life Technologies; Grand Island, NY) supplemented by 10% fetal bovine serum, 1% penicillin streptomycin, and nonessential amino acids. All cell cultures were maintained at 37°C and 5% CO_2_.

The rescued mEFs (expressing vimentin) were created by PCR amplification of the vimentin coding sequence using CloneAmp polymerase (Clontech) from pcDNA4- vimentin (provided by J. Eriksson) using the primers ggcgccggccggatccATGTCCACCAGGTCCGTGTCC and actgtgctggcgaaTTATTCAAGGTCATCGTGATGCTGAG ^57^. The PCR product was purified from an agarose gel and inserted into pBABE-hygro (pBABE-hygro was a gift from Hartmut Land & Jay Morgenstern & Bob Weinberg (Addgene plasmid # 1765; http://n2t.net/addgene:1765; RRID:Addgene_1765) cut with BamHI and EcoRI using In-Fusion (Clontech). Virus was produced by transfection of 293FT cells with pBABE- vimentin and pCL-Eco using Xfect transfection reagent (Clontech) and collection of supernatants 48 and 72hrs post transfection. The pooled virus supernatants were diluted in fresh complete medium and brought to 8ug/ml polybrene prior to addition to vim −/− mEFs. The virus supernatant was removed after 6hrs and replaced with fresh medium. 24hrs after the first application of virus supernatant, the process was repeated. 48hrs following the second application of virus, the medium was replaced with fresh complete medium containing 200ug/ml hygromycin. The selection medium was changed every two days for 7 days with the culture passaged as needed.

### 2. Immunofluorescence and microscopy

Cells were fixed either in methanol for 10 min at −20°C; or in paraformaldehyde for 30 min followed by permeabilization in 0.05% Triton X-100 in PBS for 15 min at room temperature (RT). Following fixation cells were exposed to 1% bovine serum albumin for 30 min at RT. Fixed cells were stained at room temperature for 1 hr at RT or overnight at 4° C for single or double label immunofluorescence with the following primary antibodies: chicken anti-vimentin (1:200, Novus NB300-223); rabbit anti-Lamin A (#323 from the Goldman lab); mouse monoclonal anti-Lamin B1/2 (2B2 from the Goldman lab) at a dilution of 1:1000). DNA damage repair foci were stained with monoclonal mouse anti-ƔH2AX at (1:500, EMD Millipore Corp, USA, 05-636) overnight at 4°C. Following fixation, cells were washed in PBS at RT and then incubated with secondary antibodies. This included goat anti-chicken Alexa Fluor 488 (1:1000, Invitrogen A-11039), goat anti-rabbit Alexa Fluor 488 (1:1000, Thermo Fisher A-11008), goat anti-mouse Alexa Fluor 568 (1:1000, Thermo Fisher A-11004), and goat anti-mouse Alexa Fluor 488 (1:500, Thermo Fischer A- 11029). Cells were washed and stained with Hoechst 33342 (Molecular probes H- 1399) and in some cases Alexa Fluor 647 phalloidin (Molecular Probes, 8940S) for 1 hr according to manufacturer’s instructions.

For the NLS-GFP experiments (Fig. 4) and necrosis assays (Fig. 6), cells were imaged with an inverted Leica DMIRE2 microscope equipped with either a 10x / 0.3 NA 40x/0.55 NA air objective. For the Transwell migration assays (Fig. 5 and 6), cells were imaged with an inverted Leica TCS SP8 confocal microscope equipped with a 40x/1.2 NA water objective.

For the VIF cage / F-actin visualization, 3D measurements, and sphericity analysis experiments, A Zeiss 510 LSM inverted confocal microscope with a 63x (oil immersion, NA=1.4) objective lens and excitation sources of Ar (458, 488, 514 nm), HeNe (543 nm), HeNe (633 nm) were used to create confocal stacks (Carl Zeiss, Thornwood, NY). For SIM images, the coverslips containing fix-immunostained cells were mounted on slides using Prolong Diamond (Thermo Fischer Scientific, Grand Island, MA). 3D-SIM was carried out with a Nikon Structured Illumination Super Resolution Microscope System (Nikon N-SIM; Nikon, Tokyo, Japan) with an oil immersion objective lens CFI SR (Apochromat TIRF100×, 1.49 NA; Nikon).

### 3. Immunoblotting

Lysates in 1x SDS sample buffer containing equal cell numbers of vim +/+ and vim −/− mEFs were run in triplicate on an SDS-PAGE gel and electrophoretically transferred to nitrocellulose. After blocking in 5% non-fat dry milk in PBS with 0.1% Tween 20, the blot was probed with 1:4000 dilution of chicken anti-vimentin antibody (in blocking buffer overnight at 4°C). After washing 3X for 5 mins each in PBS with 0.1% Tween 20, the blot was probed with 1:15,000 dilution of IRDye 800CW Donkey anti-chicken IgG (LI-COR) in 5% non-fat dry milk in PBS with 0.2% Tween 20 for 45 mins at room temperature. After washing 3X for 5 mins each in PBS with 0.2% Tween 20, the blot was allowed to air dry in the dark. Imaging was performed on a LI-COR Odyssey Fc and the resulting image was analyzed using LI-COR image Studio version 5.2. The amount of vimentin in each type of cell is expressed as an average of the three replicates.

### 4. Three-dimensional (3D) collagen gel preparation and imaging

Collagen gels (2 mg/mL) were prepared with final concentrations of 400,000 cells/mL. The procedure involved mixing in order pelleted and counted cells in culture medium (10% v/v), 5x DMEM (20% v/v), FBS (10% v/v), 0.1 M NaOH (10% v/v), and 4 mg/mL collagen type 1 (Corning, REF 354236, 50 %v/v). Reagents are kept cold on ice while mixing. One ml of mixture was added to 20 mm dishes and maintained at 37°C and 5% CO_2_. Experiments are conducted 24 hr after seeding the cells in the gel. The nuclei of vim +/+ and vim −/− mEFs were fluorescently labeled with by transient transfection with pEGFP-C1-NLS, 48 hr prior to seeding in collagen gel. Cell nuclei were imaged at 10 min increments for 18 hr by using wide field fluorescence with a 10x, 0.3 NA objective.

### 5. Fourier transformation

To quantify the structural features of nuclei, we traced the contour r(θ) of lamin A stained nuclei. The images were captured from methanol fixed cells with the Leica TCS SP8 confocal microscope equipped with a 40x water immersion lens. Each nucleus was traced manually and its contour interpolated from 0 to 2π by 200 points. Next, the shape fluctuations were calculated as u(θ) = r(θ) − *r*_0_, where *r*_0_ is the average radius. Next, the wave number dependent Fourier modes of the fluctuations *u*_*q*_, were obtained as

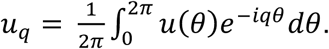

Here, the wave number *q* varies from approximately 0.15 to 30, which corresponds to 0.01 π and 2π. The average shape fluctuations were quantified by computing the Fourier mode magnitude square *u*^2^_q_ and averaging over 25 cells from two experiments.

### 6. Transwell migration assays

Cells Vim+/+ and vim −/−mEFs were seeded at sub-confluent concentrations (10-20 thousand cells/cm^2^) on polycarbonate Transwell membranes with pore diameters of either 3, 5, or 8 μm (Corning). Membranes were pre-coated with collagen 1 (50 ug/mL). mEF-seeded filters were maintained at 37°C and 5% CO_2_ for 18 hours. The cells were then gently removed from either the top or bottom of the membrane with a cotton swab and immediately fixed with either paraformaldehyde or methanol, stained, and imaged. Nuclear morphologies were determined by Hoechst staining and manually tracing nuclei using ImageJ (National Institutes of Health). Bi-nucleated cells and nuclei less than 75 μm^2^, which were likelyto be micro-nuclei, were excluded. Experiments were conducted a minimum of two times and a total of *90*-100 cells were imaged and analyzed on the top of the membrane and 65-80 cells on the bottom for both the vim +/+ and vim −/− mEFs. To determine the rate of migration through the pores, the cells were stained with crystal violet after fixation, imaged at multiple locations across the membrane with a 10x objective. Cells were manually counted in 800×800 μm^2^ fields of view (12-30 locations per condition). The percentage of cells that cross the membrane was then determined by the ratio of the number of cells on the bottom of the membrane and the sum of cells on the filter top and bottom.

### 7. Axial compression of cell-embedded collagen matrices

Collagen gels embedded with either vim +/+ or vim −/− mEFs (400 000 cells/mL; see above) were prepared in petri dishes such that the dimensions of the gel were 20 mm in diameter and 1 mm in height. After 24 hours, the gels were deposited onto a Kinexus rheometer (Malvern), equipped with a 20 mm circular parallel plate geometry and Peltier plate that maintains the sample temperature at 37 °C. The upper plate was lowered to contact the gel. Cell culture medium is added around the sample to prevent drying and allow free fluid flow. The gel is allowed to relax for a minimum of 3 minutes between the plates, then subjected to a step compression of 25%, 50% or 80% strain, which is held for 3 minutes to allow for normal stresses to relax. Then, the plate is raised, and the gel is transferred to a plate with cell culture media. Cell fate was accessed in 3D collagen gels by staining with propidium iodide (Invitrogen, REF V13241) according to the manufacturer’s instructions. Propidium iodide is normally impermeable to the cell membrane and positive staining indicates rupture of the cell membrane and necrotic cell death. Then, cells were imaged at multiple locations throughout the gel with a 10x objective and manually counted in 800×800 μm^2^ areas (5-8 locations per condition). Experiments were conducted a minimum of two times, yielding 300-700 cells per condition.

### 8. Statistical Analysis

Data presented as mean values ± standard errors (SE). Each experiment was performed a minimum of two times unless otherwise stated. The unpaired Student’s t-test with two tails at the 95% confidence interval was used to determine statistical significance. Denotations: *, p <= 0.05; **, p < 0.01; ***, p < 0.001; ns, p > 0.05. The Fisher’s exact test was used to confirm statistical significance between the proportion of blebbed and ruptured nuclei (Fig. 1) and proportion of necrotic cells (Fig. 6b) in vim +/+ and vim −/− mEFs.

### 9. 3D measurements and sphericity analysis

Confocal Z-stacks (200nm optical sections) of the cell/nucleus from vim +/+ (n=100), vim −/− (n=135), and rescued (n=25) mEFs were imported into Imaris v9.2 software to generate 3D renderings as described previously ^58^. Alexa Fluor™647 Phalloidin and Lamin A staining were used to demarcate the boundaries of the cell and the nucleus. Cell volume, nucleus volume, and nucleus surface area were calculated using the 3D renderings. Nuclear sphericity is defined as π⅓(6*V*^2/3^)/*A*, where V is the nuclear volume and A is the surface area. A value of 1 corresponds to a sphere with non-spherical shapes <1. When determining the slope of the cell versus nuclear volume in Figure 3e, cells that corresponded to two-standard deviations away from the linear regression analysis (< 5% of the cells) were defined as outliers and hence, were excluded from the analysis.

### 10. Atomic force microscopy

AFM measurements were made using a BioScope Resolve (Bruker, Santa Barbara, CA) coupled to an inverted fluorescence microscope with 10x (NA=0.3) and 20x (NA=0.8) objective lenses (Carl Zeiss, Thornwood, NY). Rounded AFM probes (10µm spheres mounted on silicon nitride cantilevers with a nominal spring constant of 0.01 N/m, Novascan Technologies, Ames, Iowa) were used. Indentation depths were kept below 500 nm to avoid any substrate effect (Rico et al. 2005) and measurements were made above the cell nucleus. Young’s modulus was determined using a modified Hertzian analysis ^39^. AFM measurements were made on mEFs grown on glass coverslips in culture medium containing 10% fetal bovine serum. At least, twenty measurements were made for each cell type.

## Acknowledgements

This work was supported by the National Institutes of Health-National Institute of General Medical Sciences (P01 GM096971). We thank Dennis Discher and Manu Tewari for help with plasmid amplification. Supported by NIGMS 2 P01 GM096971 awarded to R.D. Goldman and P. Janmey. A.V. is supported by an NCI training fellowship (T32 CA080621-15).

## Author Contributions

A.E.P. and A.V. contributed equally to this work. A.E.P. designed, performed, and analyzed experiments involving nuclear membrane rupture, collagen gels, and constricted pore migration. A.V. designed and carried out the nuclear shape and volume measurements and the AFM analyses. S.A. provided the constructs used for all analyses and carried out the immunoblotting assays. A.E.G did the SIM imaging. A.E.P, A.V., S.A., R.D.G., and P.A.J. contributed to project design and manuscript preparation.

## Supplementary Information

**SI Figure 1:**
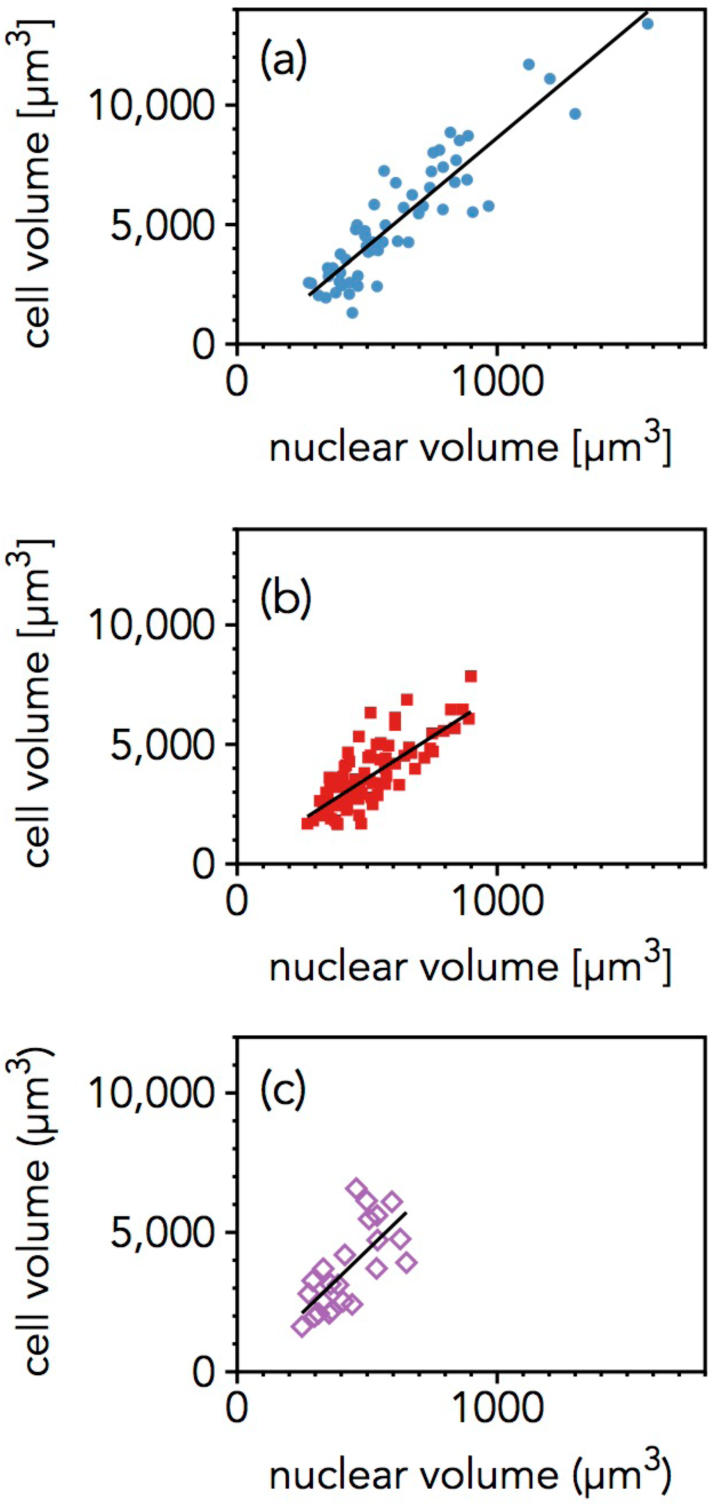
Cell volume versus nuclear volume measurements for vim +/+ (a), vim −/− (b), and rescued mEFs (c), with linear regression slopes of 9.5, 6.9, and 9.0, respectively.

**SI Figure 2:**
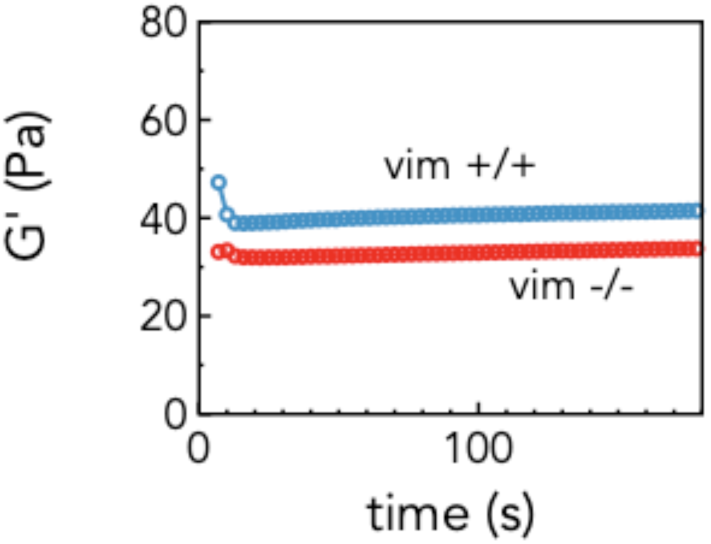
The shear stiffness of cell-embedded collagen networks in a parallel-plate rheometer. The test is performed at 2% strain amplitude and frequency 1 Hz. In the uncompressed state, there was no significance between the stiffness of the wild-type and vimentin-null mEF embedded gels (n = 4+ per condition).

**SI Figure 3:**
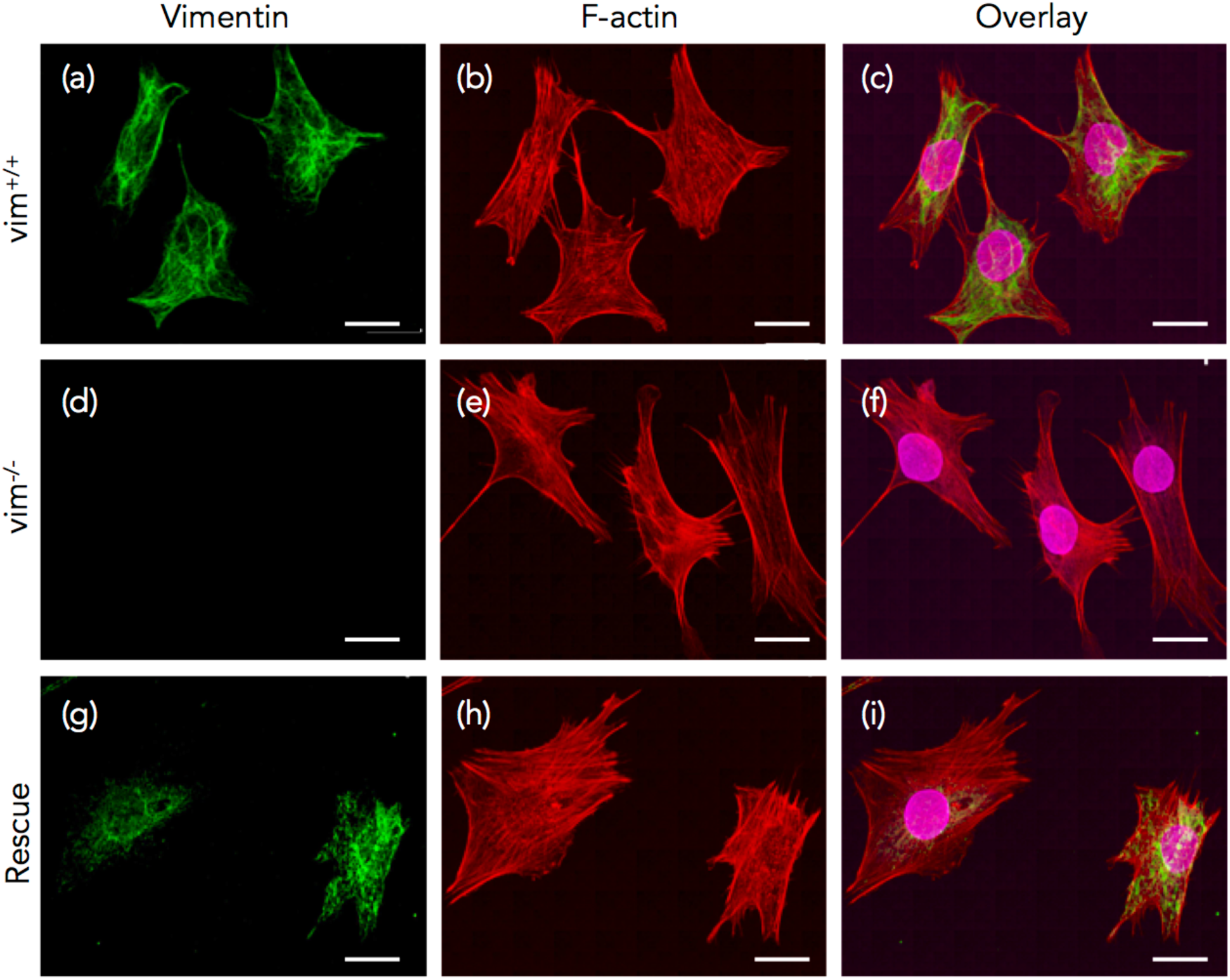
Immunostaining images of vimentin (green), F-actin (red), and lamin A (magenta) in vim +/+ (a-c), vim −/− (d-f), and rescued (g-i) mEFs expressing exogenous wild-type vimentin. Images correspond to the confocal images in Figure 2 in the main text. Scale bar is 20 μm.

### SI Movies

**SI Movie 1:** Spontaneous nuclear envelope rupture of vim −/− mEF on tissue culture plastic. The length of the video is 2.3 hr. Images are taken at 2-minute increments with a 40x air objective. Scale bar is 20 μm.

**SI Movie 2:** Spontaneous nuclear envelope rupture of vim −/− mEF in 3D collagen gel. The length of the video is 11.5 hr. Images are taken at 10-minute increments with a 10x air objective. Scale bar is 20 μm.

## References

1. Yang, J. & Weinberg, R. A. Epithelial-Mesenchymal Transition: At the Crossroads of Development and Tumor Metastasis. Developmental Cell 14, 818–829 (2008).

2. Thiery, J. P., Acloque, H., Huang, R. Y. J. & Nieto, M. A. Epithelial-Mesenchymal Transitions in Development and Disease. Cell 139, 871–890 (2009).

3. Hay, E. D. The mesenchymal cell, its role in the embryo, and the remarkable signaling mechanisms that create it. Dev. Dyn. 233, 706–720 (2005).

4. Mendez, M. G., Kojima, S. I. & Goldman, R. D. Vimentin induces changes in cell shape, motility, and adhesion during the epithelial to mesenchymal transition. The FASEB Journal 24, 1838–1851 (2010).

5. Satelli, A. & Li, S. Vimentin in cancer and its potential as a molecular target for cancer therapy. Cell. Mol. Life Sci. 68, 3033–3046 (2011).

6. Lowery, J., Kuczmarski, E. R., Herrmann, H. & Goldman, R. D. Intermediate Filaments Play a Pivotal Role in Regulating Cell Architecture and Function. J. Biol. Chem. 290, 17145–17153 (2015).

7. Dupin, I., Sakamoto, Y. & Etienne-Manneville, S. Cytoplasmic intermediate filaments mediate actin-driven positioning of the nucleus. Journal of Cell Science 124, 865–872 (2011).

8. Neelam, S. et al. Direct force probe reveals the mechanics of nuclear homeostasis in the mammalian cell. PNAS 112, 5720–5725 (2015).

9. Guo, M. et al. The Role of Vimentin Intermediate Filaments in Cortical and Cytoplasmic Mechanics. Biophysical Journal 105, 1562–1568 (2013).

10. Nekrasova, O. E. et al. Vimentin intermediate filaments modulate the motility of mitochondria. Molecular biology of the cell 22, 2282–2289 (2011).

11. Murray, M. E., Mendez, M. G. & Janmey, P. A. Substrate stiffness regulates solubility of cellular vimentin. Molecular biology of the cell 25, 87–94 (2013).

12. Ketema, M., Kreft, M., Secades, P., Janssen, H. & Sonnenberg, A. Nesprin-3 connects plectin and vimentin to the nuclear envelope of Sertoli cells but is not required for Sertoli cell function in spermatogenesis. Molecular biology of the cell 24, 2454–2466 (2013).

13. Burke, B. & Stewart, C. L. Functional Architecture of the Cell’s Nucleus in Development, Aging, and Disease. Mouse Models of The Nuclear Envelopathies and Related Diseases 109, 1–52 (Elsevier Inc., 2014).

14. Dechat, T., Adam, S. A., Taimen, P., Shimi, T. & Goldman, R. D. Nuclear Lamins. Cold Spring Harbor Perspectives in Biology 2, a000547–a000547 (2010).

15. Lammerding, J. et al. Lamins A and C but Not Lamin B1 Regulate Nuclear Mechanics. J. Biol. Chem. 281, 25768–25780 (2006).

16. Stephens, A. D., Banigan, E. J., Adam, S. A., Goldman, R. D. & Marko, J. F. Chromatin and lamin A determine two different mechanical response regimes of the cell nucleus. Molecular biology of the cell 28, 1984–1996 (2017).

17. Dahl, K. N., Engler, A. J., Pajerowski, J. D. & Discher, D. E. Power-Law Rheology of Isolated Nuclei with Deformation Mapping of Nuclear Substructures. Biophysical Journal 89, 2855–2864 (2005).

18. Swift, J. et al. Nuclear Lamin-A Scales with Tissue Stiffness and Enhances Matrix-Directed Differentiation. Science 341, 1240104–1240104 (2013).

19. Broers, J. L. V. et al. Decreased mechanical stiffness in LMNA−/− cells is caused by defective nucleo-cytoskeletal integrity: implications for the development of laminopathies. Human Molecular Genetics 13, 2567–2580 (2004).

20. De Vos, W. H. et al. Repetitive disruptions of the nuclear envelope invoke temporary loss of cellular compartmentalization in laminopathies. Human Molecular Genetics 20, 4175–4186 (2011).

21. Shimi, T. et al. The A- and B-type nuclear lamin networks: microdomains involved in chromatin organization and transcription. Genes & Development 22, 3409–3421 (2008).

22. Denais, C. et al. Nuclear envelope rupture and repair during cancer cell migration. Science 352, 353–358 (2016).

23. Raab, M. et al. ESCRT III repairs nuclear envelope ruptures during cell migration to limit DNA damage and cell death. Science 352, 359–362 (2016).

24. Irianto, J. et al. DNA Damage Follows Repair Factor Depletion and Portends Genome Variation in Cancer Cells after Pore Migration. Current Biology 27, 210–223 (2017).

25. Janmey, P. A., Euteneuer, U., Traub, P. & Schliwa, M. Viscoelastic Properties of Vimentin Compared with Other Filamentous Biopolymer Networks. Journal of Cell Biology 113, 155–160 (1991).

26. Kreplak, L., Bär, H., Leterrier, J. F., Herrmann, H. & Aebi, U. Exploring the Mechanical Behavior of Single Intermediate Filaments. Journal of Molecular Biology 354, 569–577 (2005).

27. Sarria, A., Lieber, J. G., Nordeen, S. K. & Evans, R. M. The presence or absence of a vimentin-type intermediate filament network affects the shape of the nucleus in human SW-13 cells. Journal of Cell Science 107, 1593–1607 (1994).

28. Sackmann, E. Membrane bending energy concept of vesicle- and cell-shapes and shape-transitions. FEBS Letters 346, 3–16 (1994).

29. Coordinated increase of nuclear tension and lamin-A with matrix stiffness outcompetes lamin-B receptor that favors soft tissue phenotypes. 1–28 (2017). doi:10.1091/mbc.E17-06-0393)

30. Vargas, J. D., Hatch, E. M., Anderson, D. J. & Hetzer, M. W. Transient nuclear envelope rupturing during interphase in human cancer cells. Nucleus 3, 88–100 (2012).

31. Eckes, B. et al. Impaired mechanical stability, migration and contractile capacity in vimentin- deficient fibroblasts. J Cell Sci 111, 1897–1807 (1998).

32. Helfand, B. T. et al. Vimentin organization modulates the formation of lamellipodia. Molecular biology of the cell 22, 1274–1289 (2011).

33. Friedl, P. & Bröker, E.-B. T Cell Migration in Three-dimensional Extracellular Matrix: Guidance by Polarity and Sensations. Developmental Immunology 7, 249–266 (2000).

34. Davidson, P. M., Denais, C., Bakshi, M. C. & Lammerding, J. Nuclear Deformability Constitutes a Rate-Limiting Step During Cell Migration in 3-D Environments. Cel. Mol. Bioeng. 7, 293–306 (2014).

35. Harada, T. et al. Nuclear lamin stiffness is a barrier to 3D migration, but softness can limit survival. J Cell Biol 204, 669–682 (2014).

36. Bennett, R. R. et al. Elastic-Fluid Model for DNA Damage and Mutation from Nuclear Fluid Segregation Due to Cell Migration. Biophysical Journal 112, 2271–2279 (2017).

37. Nakamura, A. J., Rao, V. A., Pommier, Y. & Bonner, W. M. The complexity of phosphorylated H2AX foci formation and DNA repair assembly at DNA double-strand breaks. Cell Cycle 9, 389–397 (2014).

38. Mendez, M. G., Restle, D. & Janmey, P. A. Vimentin Enhances Cell Elastic Behavior and Protects against Compressive Stress. Biophysical Journal 107, 314–323 (2014).

39. Vahabikashi, A. et al. Probe Sensitivity to Cortical versus Intracellular Cytoskeletal Network Stiffness. Biophysj 116, 518–529 (2019).

40. Ramsingh, R. et al. Cell deformation at the air-liquid interface induces Ca 2+-dependent ATP release from lung epithelial cells. American Journal of Physiology-Lung Cellular and Molecular Physiology 300, L587–L595 (2011).

41. Conway, D. E. et al. Fluid Shear Stress on Endothelial Cells Modulates Mechanical Tension across VE-Cadherin and PECAM-1. Current Biology 23, 1024–1030 (2013).

42. Seireg, A. & Arvikar, R. J. The prediction of musclar load sharing and joint forces in the lower extremities during walking. J. Biomechanics 8, 89–102 (1975).

43. Nia, H. T. et al. Quantifying solid stress and elastic energy from excised or in situ tumors. Nat Protoc 13, 1091 EP –

44. Irianto, J., Pfeifer, C. R., Ivanovska, I. L., Swift, J. & Discher, D. E. Nuclear Lamins in Cancer. Cel. Mol. Bioeng. 1–10 (2016). doi:10.1007/s12195-016-0437-8

45. McGregor, A. L., Hsia, C.-R. & Lammerding, J. Squish and squeeze — the nucleus as a physical barrier during migration in confined environments. Current Opinion in Cell Biology 40, 32–40 (2016).

46. Dechat, T., Adam, S. A., Taimen, P., Shimi, T. & Goldman, R. D. Nuclear Lamins. Cold Spring Harbor Perspectives in Biology 2, a000547–a000547 (2010).

47. Shimi, T. et al. Structural organization of nuclear lamins A, C, B1, and B2 revealed by superresolution microscopy. Molecular biology of the cell 26, 4075–4086 (2015).

48. Stephens, A. D. et al. Chromatin histone modifications and rigidity affect nuclear morphology independent of lamins. 1–39 (2017). doi:10.1101/206367

49. Turgay, Y. et al. The molecular architecture of lamins in somatic cells. Nature 543, 261–264 (2017).

50. Block, J. et al. Nonlinear Loading-Rate-Dependent Force Response of Individual Vimentin Intermediate Filaments to Applied Strain. Phys. Rev. Lett. 118, 048101–5 (2017).

51. Kim, D.-H. et al. Volume regulation and shape bifurcation in the cell nucleus. Journal of Cell Science 129, 457–457 (2016).

52. Khatau, S. B. et al. A perinuclear actin cap regulates nuclear shape. PNAS 106, 19017–19022 (2009).

53. Wilhelmsen, K. et al. Nesprin-3, a novel outer nuclear membrane protein, associates with the cytoskeletal linker protein plectin. J Cell Biol 171, 799–810 (2005).

54. Fontao, L. et al. The interaction of plectin with actin: evidence for cross-linking of actin filaments by dimerization of the actin-binding domain of plectin. Journal of Cell Science 2065–2076 (2001).

55. Shulman, Z. et al. Generation of compartmentalized pressure by a nuclear piston governs cell motility in a 3D matrix. Science 345, 1058–1062 (2014).

56. Xia, Y. et al. Nuclear rupture at sites of high curvature compromises retention of DNA repair factors. J Cell Biol 217, 3796–3808 (2018).

57. Morgenstern, J. P. & Land, H. Advanced mammalian gene transfer: high titre retroviral vectors with multiple drug selection markers and a complementary helper-free packaging cell lines. Nucleic Acids Research 18, 3587–3596 (1990).

58. Mulholland, W. J. et al. Multiphoton High-Resolution 3D Imaging of Langerhans Cells and Keratinocytes in the Mouse Skin Model Adopted for Epidermal Powdered Immunization. J Invest Dermatol 126, 1541–1548 (2006).

